# Human MutLα activates methylpurine DNA glycosylase to induce alkylation damage cytotoxicity

**DOI:** 10.64898/2025.12.15.694403

**Authors:** Mohamed E. Ashour, Ellissa Krekeler, Monika Chandan Bhowmik, Ning Tsao, Carlos Herrera-Montávez, Miaw-Sheue Tsai, Holland Kantar, Manal Zaher, Roberto Galletto, Nima Mosammaparast

## Abstract

Alkylation chemotherapy is commonly used against tumors such as glioblastoma, yet resistance often develops through downregulation of mismatch repair (MMR). Previous work has established that loss of MMR prevents the excision of the thymine-containing strand across *O*^6^meG-T mismatches. Thus, MMR dysfunction is advantageous because it prevents a vicious cycle of attempted repair that leads to cell death. Here, we provide an alternative explanation to this prevailing mechanism of alkylation chemoresistance by MMR loss. We find that the MMR protein MutLα physically and functionally interacts with the base excision repair (BER) enzyme methylpurine DNA glycosylase (MPG), which processes common alkylation adducts, such as 7meG and 3meA. Biochemical reconstitution demonstrates that MutLα activates MPG glycosylase activity at least partly by promoting MPG substrate binding. This glycosylase stimulation requires ATP hydrolysis as well as the MLH1-interacting region on MPG. Both MutLα or its ability to interact with MPG promote the generation of alkylation-induced abasic sites in cells, which contribute to the cytotoxicity of methyl methanesulfonate (MMS), an S_N_2 alkylating agent which does not produce *O*^6^meG. Our results provide new insight into the mechanism of alkylation chemoresistance and uncover an unappreciated crosstalk between MMR and base excision repair.

## INTRODUCTION

Alkylating agents are widely used in cancer chemotherapy and function by inducing a variety of DNA base adducts, which are well-known to be repaired by specific DNA repair pathways^1,2^. Thus, hyperactivation or loss of certain repair factors involved in these pathways may lead altered sensitivity to alkylating agents. For example, loss of mismatch repair (MMR) has been shown to induce alkylation resistance. Mismatch repair (MMR) is a highly conserved DNA repair pathway that functions canonically post-replication to maintaining genome stability. Eukaryotic MMR involves MutSα (MSH2–MSH6) and MutSβ (MSH2–MSH3) recognizing single base mismatches or small insertion-deletions, respectively, while MutLα (MLH1–PMS2) directs excision and repair through its endonuclease activity^3^. However, in the context of alkylating agent-induced damage, MMR can paradoxically promote cytotoxicity. This involves *O*^6^meG, a lesion generated by S_N_1-type alkylators such the clinically utilized drug temozolomide as well as N-methyl-N′-nitro-N-nitrosoguanidine (MNNG)^4–15^. During replication, thymine is often inserted by translesion polymerases across *O*^6^meG, which is recognized by MMR. However, the action of MMR removes the T-containing strand, leading to futile cycles of MMR-induced excision and resynthesis, which in turn gives rise to persistent single-stranded DNA, ATR activation, and ultimately cell death^4–15^. Accordingly, loss of MMR provides a well-established mechanism of resistance to S_N_1 alkylating agents.

While generally sound, there are two potential logical flaws with this model. First, it remains unclear how futile MMR cycling is sustained. The misincorporation of thymidine across *O*^6^meG is not universal, with different polymerases inserting C across from *O*^6^meG ∼15-40% of the time^16–20^. Therefore, subsequent cycles of MMR activation would be expected to be reduced until it is extinguished. Second, and more strikingly, some studies have suggested that loss of MMR factors leads to resistance not only to S_N_1 alkylating agents, but also to S_N_2 alkylating agents, like methyl methanesulfonate (MMS)^21–24^. Unlike S_N_1 alkylators, S_N_2 agents are thought to produce little to no *O*^6^-linked methyl- or alkyl-adducts^1^. These findings raise the long-standing question of how MMR is activated to mediate the toxicity of S_N_2 agents and suggest that perhaps a common substrate might trigger MMR activation in response to both S_N_1 and S_N_2 agents.

This paradox may involve the base excision repair (BER) pathway, which processes the abundant alkylation adducts 3-methyladenine (3meA) and 7-methylguanine (7meG), which together comprise 85-95% of the adducts produced by both S_N_1 and S_N_2 methylating agents^25^. Upon the induction of these lesions, BER is initiated by methylpurine DNA glycosylase (MPG, also known as alkyladenine DNA glycosylase, or AAG), which recognizes and removes damaged bases to generate abasic (AP) sites^26^. These sites are subsequently processed by AP endonuclease (APE1), which cleaves the DNA backbone, allowing DNA polymerase β to insert the correct nucleotide and DNA ligase to seal the nick, restoring DNA integrity^1,27^. Under normal conditions, BER promotes cell survival by repairing these alkylated lesions. However, excessive or dysregulated BER activity can generate toxic intermediates—such as AP sites, single-strand breaks, and hyperactivation of poly(ADP-ribose) polymerase (PARP)—that compromise genome stability^1,28,29^. Indeed, the toxic potential of hyperactivated BER, in particular increased MPG activity, is well-established^30–36^.

Here, we demonstrate a different mechanism by which MMR loss causes alkylation damage resistance, which we find to involve a physical and function crosstalk between MMR and BER. We find that the MMR protein MutLα physically interacts with MPG. This physical interaction, as well as the ATPase activity of MutLα, stimulates the glycosylase activity of MPG. Importantly, the ability of MPG to interact with MutLα, as well MutLα itself, are both critical for the generation of AP sites upon induction of alkylation. Our work provides an alternative mechanistic explanation for how MMR loss may induce chemoresistance to base damaging agents.

## Results

### Mismatch repair contributes to alkylation chemosensitivity independent of O^6^meG

We analyzed recently published genome-wide CRISPR/Cas9 screens^24^ for factors critical for mediating resistance to the commonly used alkylating agents MNNG and MMS. These results revealed that, for either drug, loss of MutLα (MLH1 and PMS2) or MutSα (MSH2 and MSH6) resulted in a significant resistance specifically to alkylation but not the other types of DNA damage (**Figure 1a-b** and **Supplemental Figure S1a**). For MNNG, this was expected, because this S_N_1 alkylating agent induces a significant amount of *O*^6^meG, which activates mismatch repair as described earlier^4–17^. However, for MMS, the screen showed nearly identical results despite the fact that this S_N_2 alkylator has been reported to produce relatively little *O*^6^meG^1^. We validated these results by showing that loss of MLH1 in either RPE-1 or HeLa cells resulted in increased resistance to MMS (**Figure 1c-d and Supplemental Figure S1b-d**). Re-expression of MLH1 rescued the sensitivity to MMS in Hela cells (**Figure 1e** and **Supplemental Figure S1e**). Using quantitative LC-MS/MS, we demonstrated that indeed, treatment of cells with MNNG creates a substantial amount of *O*^6^meG, while MMS does not (**Figure 1f**).

**Figure 1.**
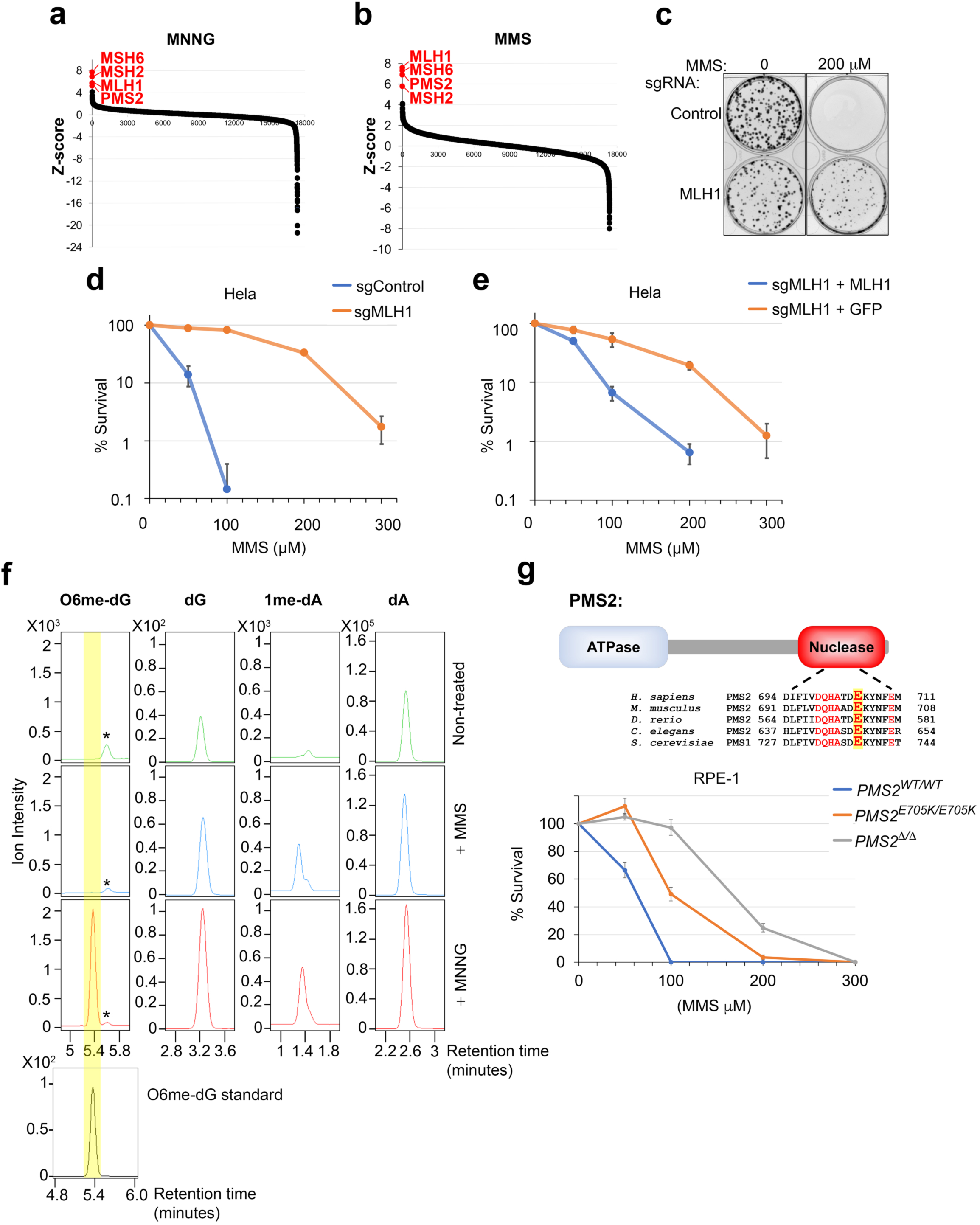
Mismatch repair promotes alkylation damage sensitivity independently of *O*^6^meG. **(a–b)** Data mining of published CRISPR/Cas9 screening datasets in RPE-1 cells treated with MMS or MNNG^24^, highlighting canonical MMR genes implicated in the resistance to alkylating damage. **(c)** Representative plate image for colony formation assay comparing isogenic wildtype (control) versus MLH1 knockout (KO) HeLa cells in response to MMS. **(d)** Colony formation assays comparing control and MLH1 knockout HeLa cells following MMS treatment. Data represent the mean of three independent biological replicates, with error bars representing ±SD of the mean; *p* < 0.0001 by two-way ANOVA. **(e)** Colony formation assay results as in **(d)** comparing MLH1-KO Hela cells complemented with Flag-GFP or Flag-MLH1 as indicated; *p* < 0.01 by two-way ANOVA. **(f)** Quantitative LC-MS/MS analysis of *O*^6^methylguanine (*O*^6^mG) and 1-methyladenine nucleoside adduct formation in RPE-1 cells following treatment with MMS or MNNG (1mM each for one hour). The *O*^6^methylguanine nucleoside standard is shown below. Asterisks indicate a nonspecific methylguanine adduct present in all biological samples. Results are representative of three independent replicates. **(g)** Colony formation assays of MMS-treated RPE-1 cells comparing WT, PMS2 knockout, and homozygous endonuclease-deficient PMS2 (PMS2*^E705K/E705K^*), illustrating the partial requirement of PMS2 nuclease activity for the MMS response; overall p value <0.0001 by two-way ANOVA.

These data suggest that MMR plays a role in alkylation damage resistance beyond its activation through induction of *O*^6^meG. Canonical mismatch repair requires the endonuclease activity of MutLα, which is encoded within the C-terminal domain of PMS2^37^. To test whether this enzymatic activity plays a role in alkylation damage resistance, we created a cell line encoding a homozygous, endonuclease-deficient PMS2 (*PMS2^E705K/E705K^*) as well as a knockout (*PMS2^Δ/Δ^*; **Figure 1g** and **Supplemental Figure S1f-g**). While PMS2 loss led to MMS resistance, the *PMS2^E705K/E705K^*cells displayed an intermediate phenotype, with resistance greater than parental cells but significantly less than *PMS2^Δ/Δ^* cells (**Figure 1g**). This indicated that human MutLα has a function in alkylation damage resistance independent of canonical MMR.

### MutLα interacts physically and functionally with MPG glycosylase

To determine how MutLα may function to promote alkylation sensitivity, we performed an interactome screen of Flag-MLH1 using immunoprecipitation from HeLa-S nuclear extract followed by mass spectrometry (IP-MS; **Figure 2a** and **Supplemental Table S1**). This revealed several known MLH1 interactors, such as PMS2, EXO1, BLM and BRIP1^38–42^, as well as two DNA glycosylases, namely MBD4 and MPG. MBD4 was previously shown to interact directly with MLH1^43^. Because MPG is the key glycosylase for common alkylated lesions^1,27^, we also performed an IP-MS analysis of MPG, which revealed both components of MutLα, as well as previously identified partners, namely PCNA and the Elongator complex^44,45^ (**Supplemental Table S2**). Co-immunoprecipitation followed by Western blotting validated the interaction between MPG and MutLα components, in the presence or absence of MMS (**Figure 2b**). Using the recently developed Predictomes.org server, which uses AlphaFold Multimer to test for potential interactions between distinct pairs of genome stability proteins^46^, we found a potential direct interaction interface between MPG and MLH1 (**Figure 2c**). A closer analysis suggested that just outside of its catalytic domain, MPG contains a conserved MIP (<u>M</u>LH1 interacting <u>p</u>rotein) box in its N-terminus, a short motif shared by a number of repair factors that interact with MLH1, including EXO1 and FAN1^47–49^ (**Figure 2d-2e**).

**Figure 2.**
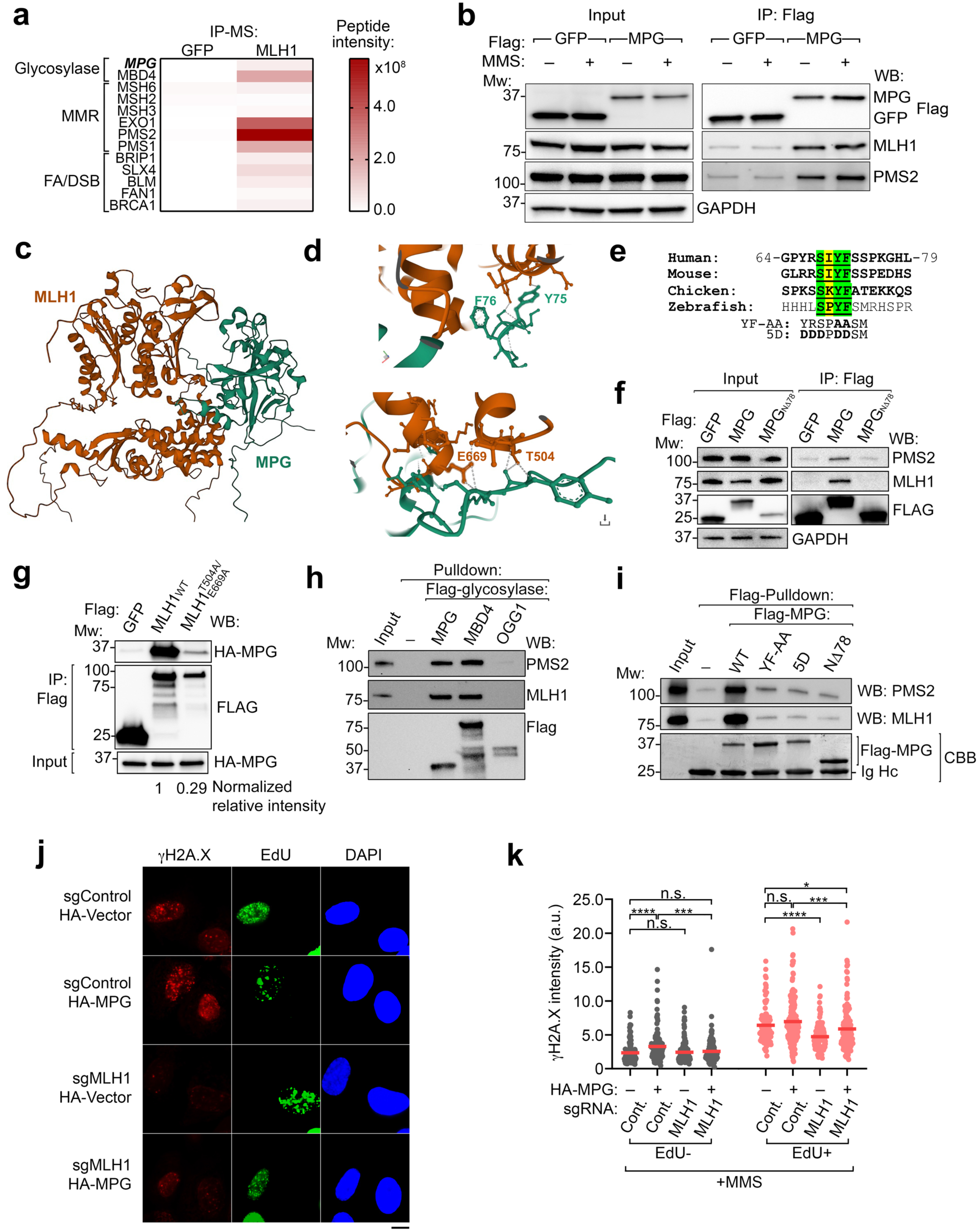
MutLα interacts both physically and functionally with MPG. **(a)** Heat map of key genome stability factors identified by immunoprecipitation followed by mass spectrometry (IP-MS) using Flag-MLH1 versus Flag-GFP. **(b)** Co-immunoprecipitation (Co-IP) of Flag-followed by western blotting validating the interaction between MPG and MutLα in with or without MMS treatment. Results are representative of three independent replicates. **(c-d)** Protein–protein interaction predictions from AlphaFold Multimer as curated by Predictomes.org^77^, indicating a potential direct association between MPG (green) and MLH1 (orange). **(e)** Sequence analysis revealing that MPG contains a conserved MLH1-interacting protein (MIP) box within its N-terminus. Targeted mutations (designated YF-AA and 5D) used in this study are shown below. Results are representative of three independent replicates. **(f)** Immunoprecipitation of Flag-GFP, Flag-MPG, and Flag-MPG^Nλ−78^ was performed using 293T cells, with input and IP material being probed with the indicated antibodies. Results are representative of three independent replicates. **(g)** Flag-GFP, Flag-MLH1 or Flag-MLH1^T504A/E669A^ were co-expressed with HA-MPG in 293T cells. After Flag-immunoprecipitation, input and IP material were probed with the indicated antibodies. Relative HA:Flag signal was quantified and is shown below. Results are representative of three independent replicates. **(h)** *In vitro* binding assays were performed using recombinant, Flag-immobilized glycosylases and untagged MutLα complex. Bound and input material were analyzed by western blot using the indicated antibodies. Results are representative of three independent replicates. **(i)** *In vitro* binding assays were performed as in **(h)** using the indicated recombinant, Flag-immobilized MPG variants and untagged MutLα. Bound and input material were analyzed by Coomassie blue (CBB) or western blot using the indicated antibodies. Results are representative of three independent replicates. **(j-k)** immunofluorescence (IF) and quantification of γH2AX foci intensity in RPE-1 cells treated with MMS, in the presence or absence of MLH1 and exogenous HA-MPG where shown, with quantification shown in **(k)**. Data represent the mean of three biological replicates. Statistical significance was determined using the Mann–Whitney test; \**p* < 0.05, \*\*\**p* < 0.001, and \*\*\*\**p* < 0.0001.

AlphaFold3 modeling using MPG and both components of MutLα gave similar results, with the predicted interaction involving MPG residues Pro70 through Phe76 contacting MLH1 residues Asn502–Cys672 and a small part of PMS2 residues spanning Gln604 to Ala608 (**Supplemental Figure S2a-b**). To test this model, we performed co-immunoprecipitation of WT MPG as well as an N-terminal deletion which removes the MIP box. Deletion of this domain (MPG NΔ78) completely abrogated MutLα interaction (**Figure 2f**). Similarly, targeted mutagenesis of MLH1 at key sites that are structurally important for MIP box recognition (Thr504Ala and Glu669Ala) also resulted in a significant, albeit incomplete, loss of co-immunoprecipitation between MPG and MutLα (**Figure 2g**). We also tested whether this interaction is direct, using recombinant His-Flag-tagged glycosylases and untagged MutLα purified from insect cells (**Supplemental Figure S2c-S2d**). Both MPG as well as MBD4, which also contains a MIP box motif^43^, bound to MutLα in these assays, while OGG1, which does not appear to have a MIP box, did not (**Figure 2h**). Consistent with these results, the direct MPG-MutLα interaction depended on the MIP box, as MPG NΔ78, or targeted point mutations within the MIP box all markedly reduced this interaction (**Figure 2e and 2i**).

Given their physical interaction, we reasoned that the processing of alkylated lesions by MPG may be functionally regulated by MutLα, which could explain the non-canonical role of MutLα in alkylation resistance. Using RPE-1 cells, we found that loss of MLH1, outside of S-phase, had little effect on MMS-induced DNA damage signaling, as measured by γH2A.X foci intensity (**Figure 2j-k and Supplemental Figure S2e**). However, in cells undergoing replication (EdU+) cells, loss of MLH1 led to a significant reduction in the MMS-induced DNA damage activation. This is consistent with MMR being activated immediately following DNA replication^8,9^. This loss of alkylation-induced DNA damage response during S-phase in MLH1-deficient cells could be at least partially reversed by expressing exogenous MPG, suggesting that the alkylated substrates for MPG are present but in the absence of MLH1 they are not processed to the same extent. In contrast to MMS-treated cells, under untreated conditions the status of MLH1 or MPG does not alter γH2A.X levels (**Supplemental Figure S2f)**. Altogether, these data suggest that a physical and functional relationship may exist between BER and MMR in the context of alkylation damage.

### MutLα directly stimulates MPG glycosylase activity

To assess the functional significance of the MutLα-MPG interaction, we performed *in vitro* glycosylase assays. Substrates containing hypoxanthine (HX), an established substrate of MPG^50,51^, were used in these assays, since alkylated lesions like 3meA and 7meG are unstable^52,53^. Interestingly, MutLα stimulated MPG activity on an HX-containing dsDNA substrate as well as on a substrate where the HX is near a ssDNA-dsDNA junction (**Figure 3a and Supplemental Figure S3a**). The stimulation was evident particularly when concentrations of MPG were limiting (**Figure 3a and Supplemental Figure S3a**). The stimulation was not due to MutLα alone having any glycosylase activity toward an HX-containing dsDNA substrate, or any AP lyase/endonuclease activity using an AP site analogue (tetrahydrofuran; THF) (**Supplemental Figure S3c-d**). However, stimulation on a ssDNA substrate was minimal (**Supplemental Figure S3b**). During replication, cytidine is preferentially incorporated across from HX, and MPG is relatively inefficient at removing HX from HX-C compared to HX-T, presumably because of the stronger base pairing properties of HX-C^54^, prompting us to test this substrate. Similar to HX-T, MutLα can also significantly stimulate MPG glycosylase activity on the HX-C substrate (**Supplemental Figure S3e**). Since MutLα possesses two ATPase domains, both of which cooperate to induce conformational changes, including in partner proteins^37,55^, we tested whether ATP hydrolysis contributes to MPG activation. While the presence of ATP promoted MutLα stimulation of MPG, the nonhydrolyzable analog AMP-PNP failed to do so (**Figure 3b**). As MBD4 also associates directly with MutLα, we tested whether MutLα promotes this particular glycosylase. However, MutLα had little impact on MBD4 activity using the well-established G:T mismatch substrate for MBD4^56^ (**Figure 3c**).

**Figure 3.**
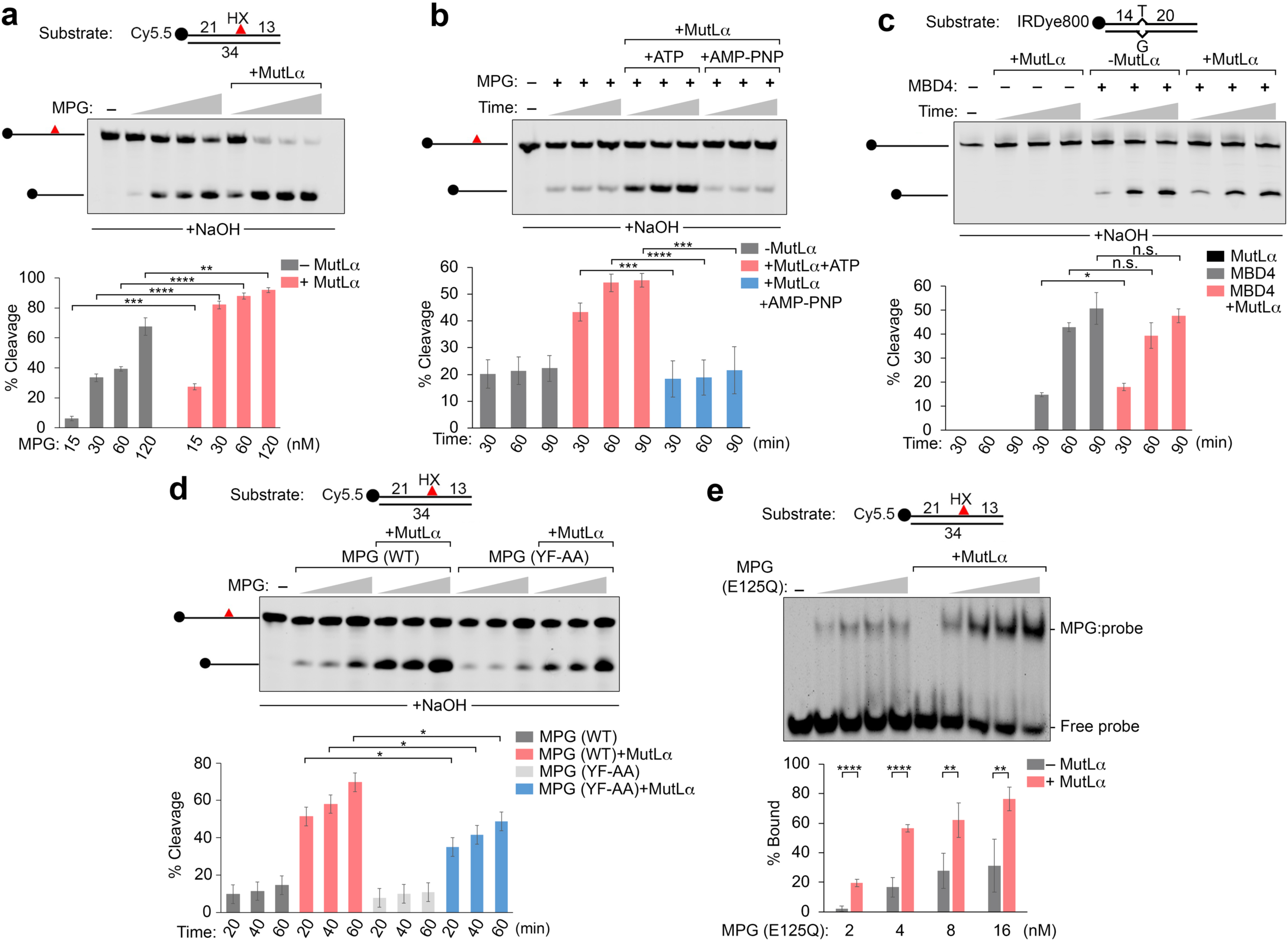
MutLα promotes MPG glycosylase activity. **(a)** *In vitro* MPG glycosylase assays using hypoxanthine (HX)-containing DNA substrates (80 nM), to examine the effect of MutLα (30 nM) on MPG glycosylase activity. **(b)** *In vitro* MPG glycosylase assays comparing the effects of ATP versus AMP-PNP on MutLα (30 nM) activation of MPG (20 nM). **(c)** *In vitro* MBD4 glycosylase assays using substrate containing a G/T mismatch (80 nM), MutLα used at 75 nM and MBD4 was used at 80 nM. **(d)** *In vitro* MPG glycosylase assays comparing wild-type (WT) MPG and the MPG MIP-box mutant (YF-AA), each at 10nM for 20-60 min, with or without MutLα (30 nM). **(e)** Electrophoretic mobility shift assays (EMSAs) using HX-containing DNA (8 nM) and a catalytically inactive MPG mutant (2-16 nM) with or without MutLα (40 nM) as indicated. Data represent the mean of at least three independent replicates ± SD of the mean. Statistical significance was determined using student’s t-test; \**p* < 0.05, \*\**p* < 0.01, \*\*\**p* < 0.001, and \*\*\*\**p* < 0.0001.

We next asked whether the direct interaction between MutLα and MPG promotes the glycosylase. We found that a MIP box mutant (MPG YF-AA) did not significantly impair MPG’s intrinsic glycosylase activity, but it modestly yet consistently reduced MutLα stimulation of MPG (**Figure 3d**). Other factors, such as APE1 and UV-DDB, stimulate MPG without direct physical interaction, but instead by competing for the abasic site product, leading to increased MPG turnover^57,58^. We found the MutLα and APE1 stimulation of MPG activity was additive (**Supplemental Figure 3f**), suggesting that MutLα may function differently from these other factors in stimulating MPG. An alternative possibility is that MPG binding to its substrate may be stimulated by MutLα. Using electrophoretic mobility shift assays (EMSAs) with an HX DNA substrate and a catalytically inactive MPG mutant^59^ (**Supplementary Figure 3g**), we found that MutLα increased binding of inactive MPG to HX-DNA (**Figure 3e**). These results indicate that MutLα stimulates MPG glycosylase activity, via ATP hydrolysis and direct physical interaction, to promote substrate binding.

### The MutLα-MPG interaction promotes alkylation cytotoxicity and abasic site accumulation

Our data suggest that the MIP box of MPG may promote its recruitment to the DNA substrate. After knock-down of endogenous MPG, quantification of MPG bound to chromatin following MMS treatment revealed that WT MPG was bound, yet the MIP box mutants were significantly reduced in this regard, even though they were expressed at similar levels (**Figure 4a-b** and **Supplemental Figure S4a-b**). To better understand the function of the MPG-MutLα interaction in cells, we quantified abasic site accumulation using an inducible HA-tagged, catalytically inactive form of APE1 endonuclease (E96Q/D210N), which has been used before for this purpose, as it retains strong binding to abasic sites^60,61^. WT MPG had greater levels of this reporter protein associated with chromatin compared to the MPG^5D^ MIP box mutant, suggesting that the MPG-MutLα interaction is important in abasic site production in cells. Although the difference between WT and MPG^YF-AA^ was not statistically significant, WT and MPG^5D^ displayed a significant difference (**Figure 4c-d** and **Supplemental Figure S4c**). Consistent with a role in promoting abasic site production, loss of MLH1 or PMS2 resulted in significantly less MMS-induced abasic site accumulation than control cells (**Figure 4e-h and Supplemental Figure S4d-S4e**). Finally, increased MPG activity has been previously established to increase alkylation cytotoxicity due to accumulation of BER intermediates, particularly abasic sites^1^. Consistent with this, expression of WT MPG significantly increased MMS-induced cytotoxicity, while the MPG^YF-AA^ mutant had a more modest effect, and the MPG^5D^ mutant behaved similar to the vector control (**Figure 4i**). Altogether, our data suggest that MutLα and specifically its interaction with MPG are important for alkylation damage responses by promoting abasic site production (**Figure 4j**).

**Figure 4.**
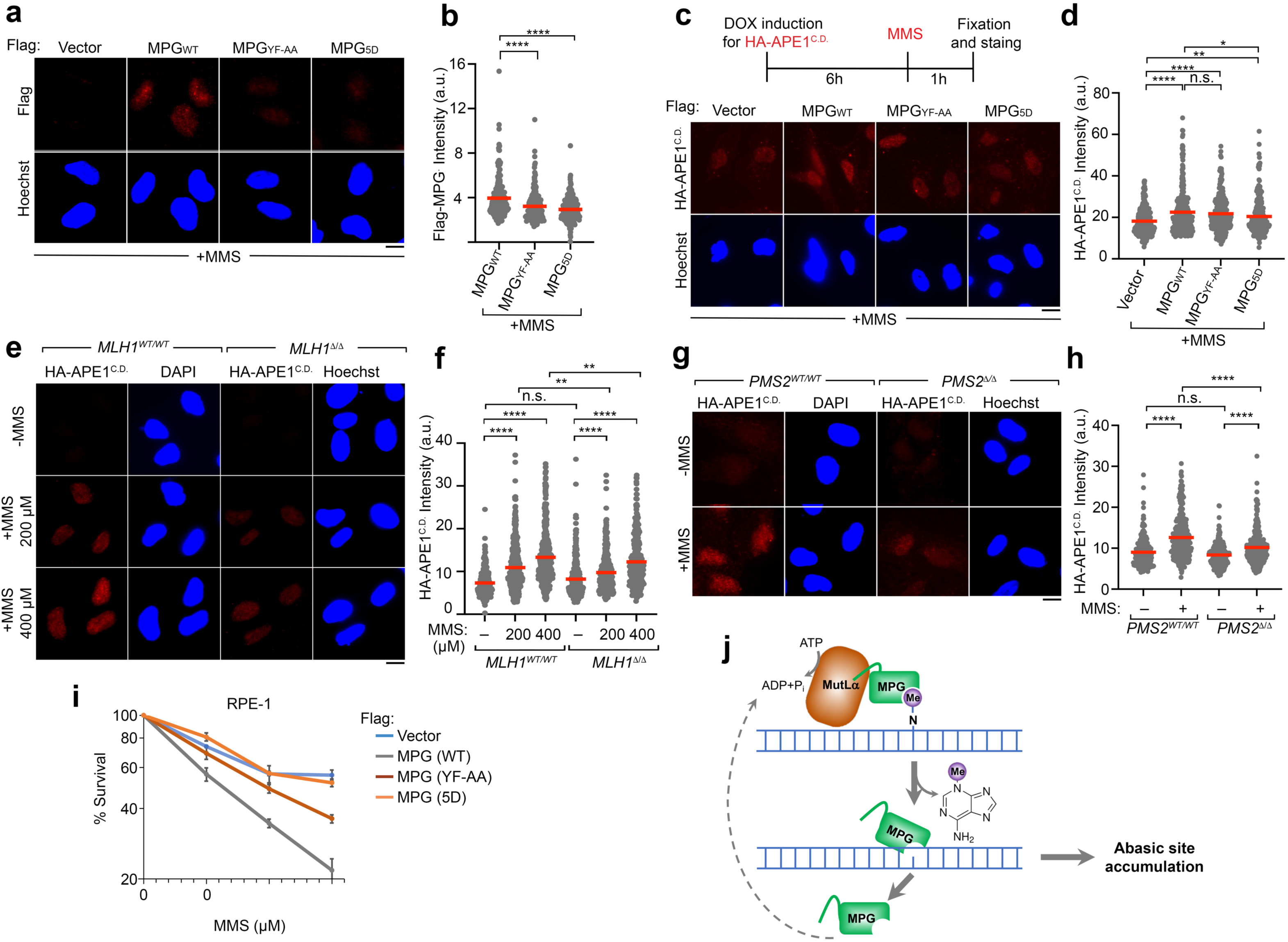
MutLα and the MPG MIP box promote abasic site accumulation and alkylation-induced cytotoxicity. **(a-b)** Immunofluorescence analysis and quantification of Flag-MPG chromatin association in RPE-1 cells treated with MMS, comparing WT MPG with MPG MIP box mutants (MPG^YF-AA^ and MPG^5D^). **(c-d)** Experimental scheme for using an HA-tagged catalytically inactive APE1 (E96Q/D210N), a reporter for abasic site accumulation. RPE-1 cells were treated with an shRNA against MPG, and the indicated Flag vectors were used for re-expression of WT MPG or MIP-box mutants. **(e-f)** Immunofluorescence analysis of abasic site accumulation as in **(a-b)** comparing control or MLH1 knockout cells with or without MMS treatment. **(g-h)** Immunofluorescence analysis and quantification of abasic site accumulation as in **(a-b)** comparing control versus PMS2 knockout cells with or without MMS treatment. Data represent the mean of at least three biological replicates. Statistical significance was determined using the Mann–Whitney test; \**p* < 0.05, \*\**p* < 0.01, and **** *p* < 0.0001. **(i)** Colony formation assays comparing EV, WT MPG, and MIP-box mutant Flag-MPG–expressing cells. Data represent the mean of three replicates, with error bars indicating ±SD of the mean. Statistical significance was assessed by two-way ANOVA; the overall P values for EV vs WT, EV vs YF/AA, and EV vs 5D were P < 0.0001, P < 0.001, and not significant, respectively. **(j)** Working model illustrating the role of MutLα in activation of MPG, releasing N-linked alkylated lesions and inducing abasic site accumulation.

## Discussion

It has been long assumed that alkylation damage resistance through loss of MMR is driven primarily through activation of this pathway upon encountering *O*^6^meG-T mismatches. Because of the removal of the undamaged strand, a futile cycle ensues, where continued MMR activation and its repair intermediates, particularly single-stranded DNA, lead to cell death ^4–15^. Here, we demonstrate an alternative mechanism for how MMR functions in response to alkylation independently of the *O*^6^meG lesion. MMR loss leads to marked alkylation damage resistance even to MMS, an S_N_2 alkylator, which produces little to no *O*^6^meG (**Figure 1**). Surprisingly, the cleavage of DNA by MutLα is partially dispensable for the increased resistance to MMS, indicating a non-enzymatic function of this nuclease in this resistance pathway. Through proteomic analysis and AlphaFold-guided modeling, we identified MPG as a direct partner of the MutLα subunit MLH1 (**Figure 2**), and demonstrated that MutLα directly activates the glycosylase activity of MPG *in vitro* (**Figure 3**). Consistent with this, we show that MutLα, as well as the MLH1-interaction surface on MPG, are both important for abasic site generation in cells, as well as alkylation damage responses (**Figure 4**). To our knowledge, this represents the first direct mechanistic link between the MMR and the generation of abasic sites in cells, revealing a previously unrecognized role for MMR in promoting BER-associated intermediates.

Together, our data support a model where two distinct pathways of single-stranded DNA repair crosstalk with each other during alkylation damage (**Figure 4i**). In this model, the activation of MPG during or soon after S-phase by MutLα would create BER intermediates, in particular abasic sites, which are known to be more toxic than the initial base damage (e.g., 7meG) ^1^. Our work provides a mechanistic explanation for the observation of why MMR plays an important role for resistance to both S_N_1 alkylators, which produce a significant amount *O*^6^meG, as well as S_N_2 alkylators, which do not. It is likely that the lesions that are common to both classes of agents, such as 7meG and 3meA, both of which are processed by MPG ^1^, are responsible for the effect of MMR loss in alkylation resistance. These findings rationalize why MMR loss provides resistance across diverse classes of alkylators: it is the shared substrates of BER, rather than *O*^6^meG alone, that underlie this phenotype. We should note, however, that our model does not dismiss the futile cycle model involving *O*^6^meG – it merely expands upon it. Indeed, the catalytic activity of PMS2 is still responsible for some of the resistance to MMS, suggesting that the canonical MMR pathway involving single-stranded DNA production is still important for this resistance, albeit in a manner that is independent of *O*^6^meG (**Figure 1g**).

It is tempting to contemplate why this direct crosstalk between MutLα and MPG would evolve. Beyond improving base excision repair efficiency, it may also expand the ability of MPG to process otherwise refractory lesions, such as hypoxanthine paired with cytosine, where strong base-pairing prevents the usual base-flipping mechanism of MPG ^51,62^ Similarly, while MPG is known to bind εA and εC but efficiently cleave only εA, MLH1 may assist in processing such substrates ^63^. In contrast, under high levels of alkylation stress - such as in experimental settings or during chemotherapy - the excessive formation of abasic sites could become harmful. A second possibility is that the glycosylase activity of MPG may function to promote mammalian MMR by creating the initial nick in nascent DNA. Unlike prokaryotes which differentially mark parental versus nascent DNA through methylation, the incorporation of ribonucleotides by replicative DNA polymerases and their subsequent removal is believed to be a major mechanism by which mammalian MMR differentiates the two DNA strands. Yet loss of ribonucleotide processing impacts MMR very modestly in eukaryotes ^64,65^, suggesting other mechanisms may be at play in creating nicks in nascent DNA. Thus, activation of MPG by MutLα may serve as another way for marking the nascent strand. If this were the case, it would require a substrate for MPG to be incorporated into DNA during replication. This has been confirmed for deoxyinosine and εA, both of which are readily incorporated by DNA polymerases ^66,67^, although the amount of deoxyinosine triphosphate is kept at a minimum by inosine triphosphatase ^68,69^.

Overall, our study provides a new mechanistic framework linking MMR to BER in the response to alkylation damage, expanding on how MMR loss confers chemoresistance. This work highlights the complexity of DNA repair network crosstalk and may inform the design of therapeutic strategies that exploit MMR-BER interactions, either by targeting MPG activation or by modulating MutLα function, to enhance alkylation chemotherapy efficacy.

### Limitations of study

Despite this progress, several key questions remain. First, the precise mechanism by which MutLα stimulates glycosylase activity of MPG via ATP hydrolysis is not fully clear and requires further investigation. Second, the relative contribution of the MutLα-MPG pathway to alkylation-induced cytotoxicity, compared with canonical MMR, is not completely clear. Third, MutSα has a role in this pathway as well, since its loss causes MMS resistance, but its mechanism is unclear. It is possible that MutSα may help recruit and activate MutLα in a manner similar to canonical MMR. Yet, evidence also suggests that MutLα can function independently of MutSα and participating in processes such as meiotic recombination, double-strand break repair, and EXO1 regulation^70,71^. This raises the question of whether MPG-MutLα and MutSα−MutLα complexes act on the same or different substrates. One possibility is that MutLα partners with both recognition factors, using MutSα to detect *O*^6^meG lesions and MPG to detect 7meG and 3meA.

## Acknowledgments

We thank Peter Burgers, Hani Zaher, and Priyanka Verma for suggestions during the course of these studies, as well as Justin Leung and Priyanka Verma for providing plasmid constructs. We acknowledge Ross Tomaino (Harvard Medical School Taplin Core) for proteomics analysis, as well as the Genome Engineering and iPSC Center (GEiC) at Washington University for CRISPR engineering of cell lines. This work was supported by the National Institute of Health (R35GM139508 to R.G; R01CA282733, R01GM160854, and P01CA092584 to N.M.), an American Cancer Society Research Scholar award (RSG-18-156-01-DMC to N.M.), the Barnard Foundation and the Siteman Cancer Center (to N.M.).

## Author contributions

M.E.A. and N.M. designed the project. M.E.A., E.K., M.C.B., N.T., C.H-M., and H.K. carried out cellular and biochemical experiments, and performed data analysis with input from N.M.. M.A., M-S.T. and M.Z. performed MutLα protein purification. R.G. supervised M.Z. M.E. A. and N.M. wrote the manuscript with input from all other authors. N.M. supervised the project.

## STAR Methods

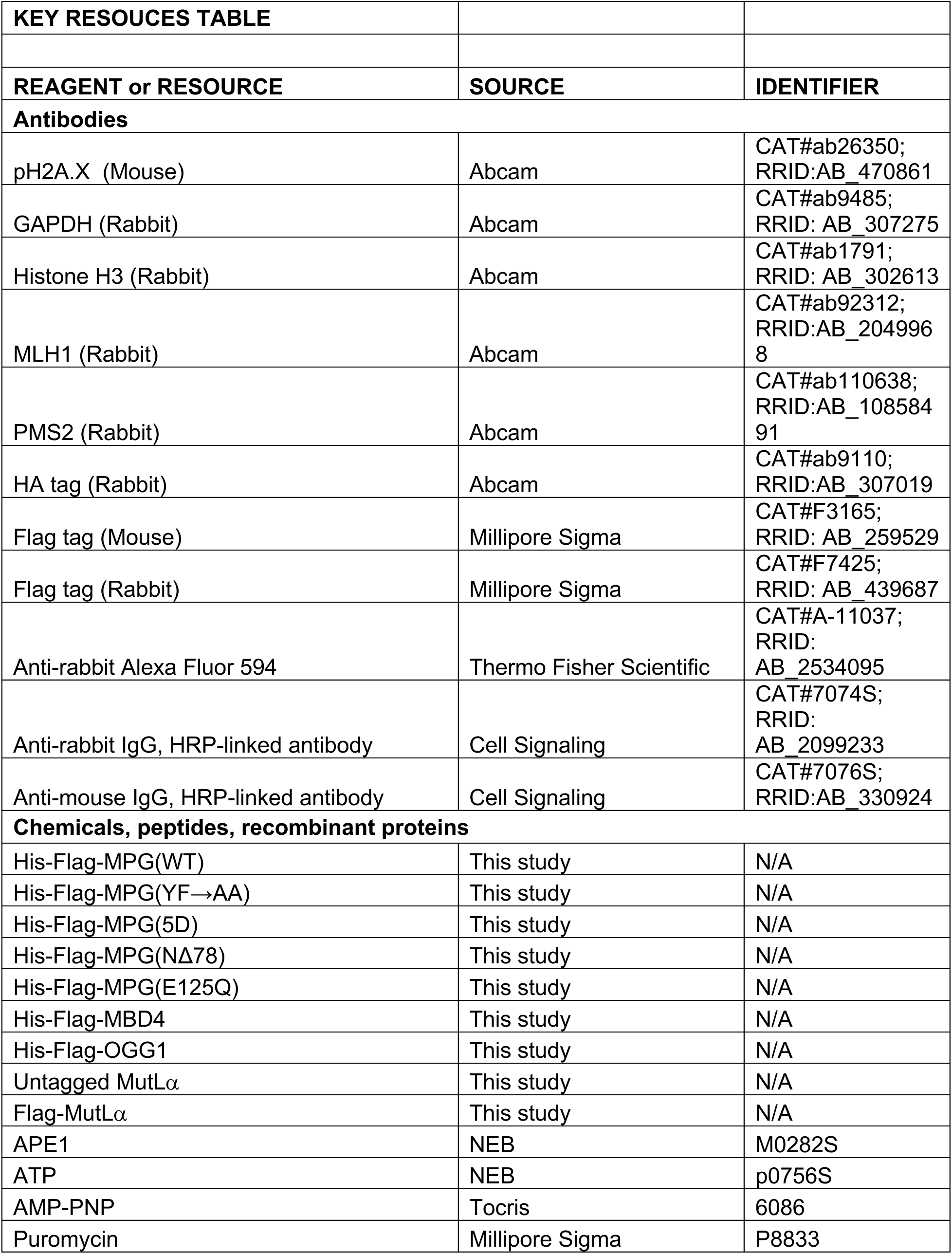
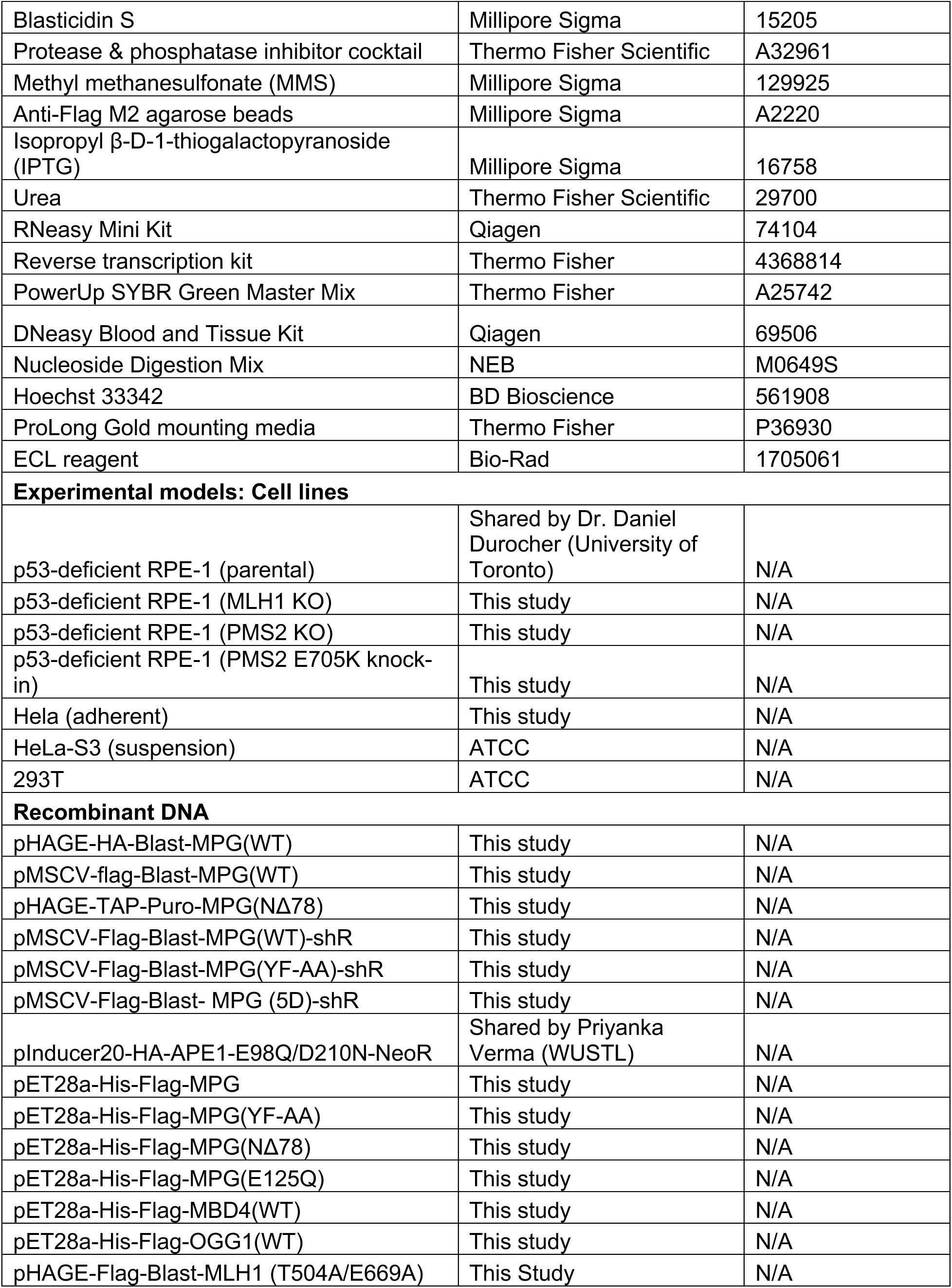
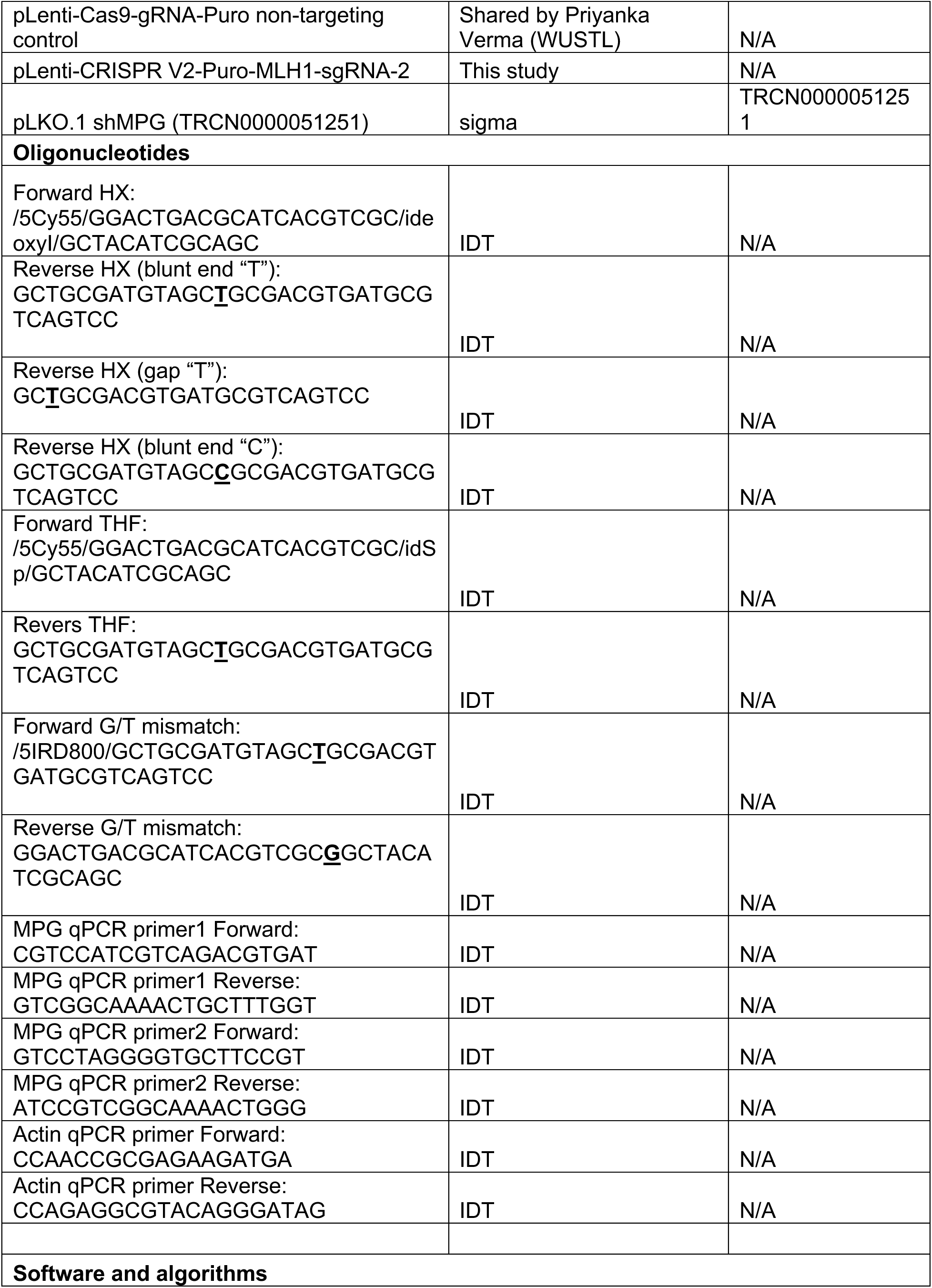
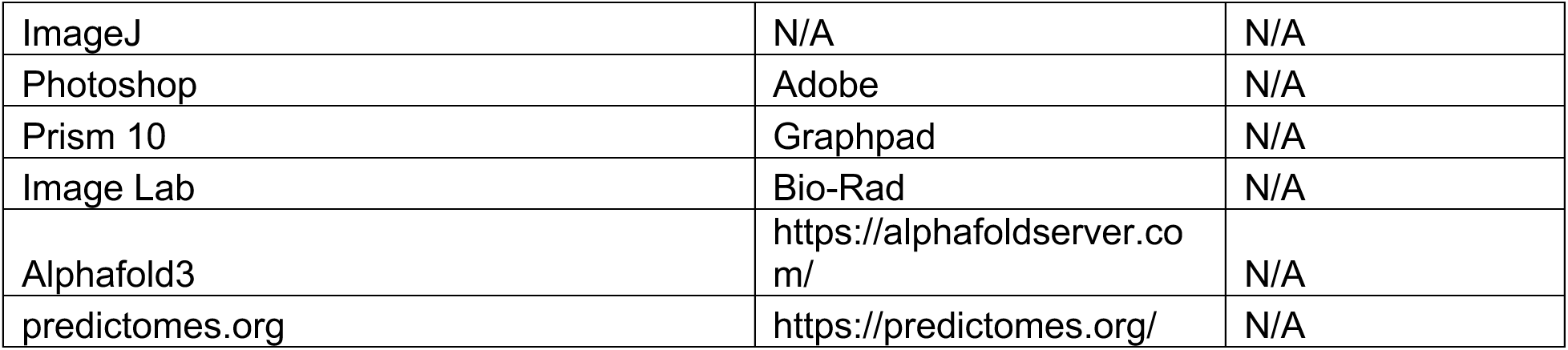

### RESOURCE AVAILABILITY

#### Lead Contact

Further information and requests for resources and reagents should be directed to and will be fulfilled by the Lead Contact, Nima Mosammaparast.

#### Materials Availability

All reagents generated in this study are available from the Lead Contact without restriction.

#### Data and Code availability

- Plasmid sequencing data will be deposited at GEO and will be made publicly available prior to publication, or upon request.
- Original images and blots will be deposited at Mendeley Data and will be publicly available as of the date of publication, or earlier upon request.
- This paper does not report original code.
- Any additional information required to reanalyze the data reported in this paper is available from the Lead Contact upon request.

### EXPERIMENTAL MODEL AND STUDY PARTICIPANT DETAILS

#### Cell culture

All human cell lines were cultured in Dulbecco’s modified eagle medium (ThermoFisher), supplemented with 10% fetal bovine serum (MilliporeSigma or Avantor Seradigm), 100 U/ml of penicillin-streptomycin (ThermoFisher) at 37°C and 5% CO_2_. Cell lines were authenticated using the ATCC human STR profiling service and regularly tested for mycoplasma contamination using PCR-based Mycoplasma Testing at the Washington University Genome Engineering and iPSC Center (GEiC). Preparation of viruses, transfection, and viral transduction were performed as described previously^72^. Briefly, replication incompetent viral particles were produced in packaging cells (HEK293T) by co-transfection with compatible packaging plasmids. Subsequently, target cell lines were transduced using supernatant from transfected HEK 293T cells, in the presence of 8 μg/ml polybrene (MilliporeSigma). Infected cells were selected with puromycin (HeLa, 1 μg/ml; RPE1-hTERT, 20 μg/ml), blasticidin (Hela, 10 μg/ml; RPE1-hTERT, 5 μg/ml) or neomycin (RPE1-hTERT at 400 μg/ml) for 48–72 h.

### METHOD DETAILS

#### Plasmids

For mammalian cell expression, MPG and MLH1 cDNA or gBlocks were subcloned into pMSCV or pHAGE-tag vector by Gateway recombination as described ^72^. For recombinant expression of MPG and MPG variants, gBlocks from IDT were cloned into pET28a-2xFlag vector using BamHI/XhoI. The coding sequences (CDS) of MBD4 and OGG1 were generously provided by Justin W. Lueng (University of Texas Health Science Center at San Antonio), and cloned by PCR into pET28a-2xFlag. All the constructs were confirmed by whole-plasmid sequencing (Plasmidsaurus). The bacterial glycerol stocks harboring sequence-verified shRNA lentiviral plasmid vectors for human genes were selected from MISSION shRNA library (MilliporeSigma). The clone used for targeting MPG was TRCN0000051251 (Target Sequence: CCTGTACGTGTACATCATTTA). Scrambled shRNA vector was used as a negative control (SHC002, from MilliporeSigma; shRNA sequence: 5′-CAACAAGATGAAGAGCACCAA-3′).

#### CRISPR/Cas9-mediated knockouts

The RPE-1 PMS2 KO and knock-in cells were created using the Genome Engineering and iPSC Center (GEiC) at Washington University, using the gRNA sequence 5’-GAAGTTATACTTCTCGTCCG 3’. PMS2 knockout/knock-in clones were verified by deep sequencing and by western blot analysis. For CRISPR/Cas9 mediated lentiviral knockout of MLH1, the gRNA sequence (5’-ACTACCCAATGCCTCAACCG-3’) gRNA sequences were cloned into pLentiCRISPR-V2 (Addgene #52961). MLH1 knockout was verified by western blot analysis.

#### MPG and MutLα complex purification and proteomic analysis

HeLa-S3 cells expressing retroviral Flag-HA-MPG or lentiviral Flag-MLH1, along with GFP-expressing controls, were harvested for nuclear extraction according to previous protocols^73^. Nuclear extracts were used for complex purification using using anti-Flag (M2) resin (Millipore Sigma) in TAP buffer (50 mM Tris-HCl pH 7.9, 100 mM KCl, 5 mM MgCl_2_, 10% glycerol, 0.1% NP-40, 1 mM DTT, and protease inhibitors). After peptide elution, the complexes were TCA precipitated and associated proteins were identified by LC-MS/MS at the Taplin Mass Spectrometry Facility (Harvard Medical School) using an LTQ Orbitrap Velos Pro ion-trap mass spectrometer (ThermoFisher) and Sequest software^74^. Total and unique peptide spectral counts, as well as total peptide intensities, can be found in **Supplemental Tables S1 and S2**.

#### Immunoprecipitation

Immunoprecipitations were performed using HEK293T cells using transient transfection, or HeLa cells using stably transduced cells as indicated in the figure legends. The cells were resuspended in high-salt buffer (50 mM Tris-HCl pH 7.9, 300 mM KCl, 10% glycerol, 1.0% Triton X-100, 1 mM DTT, and protease inhibitors), lysed by sonication, and centrifuged. An equal volume of buffer without KCl was added, and the lysate was incubated with anti-Flag (M2) resin (Millipore Sigma). After incubation at 4°C with rotation, the resin was washed extensively with buffer containing 150 mM KCl. Bound material was eluted with Laemmli buffer and analyzed by Western blotting.

#### Recombinant protein purification

For His-tagged glycosylases, Rosetta (DE3) bacteria were transformed with pET-28a-based vectors and grown in LB broth plus Kanamycin. After initial growth at 37°C, cells were induced with 0.5 mM IPTG and grown at 20°C overnight. Protein purification was performed via FPLC on an ATKA GO using a 1 mL HisTrap HP column (Cytiva). Proteins were eluted from the column on a gradient reaching 500mM Imidazole. Peak fractions were pooled and dialyzed overnight using TAP wash buffer.

Untagged hMutLα purification was performed with modifications to the previously described method^37^. Sf9 cells were continuously cultured in 800 ml of serum-free HyQ-SFX medium (HyClone, Inc.) at 27 °C in 2.8-liter Fernbach flasks. When culture density reached 1 × 10^6^/ml, cells were co-infected with hMLH1 and hPMS2 baculovirus constructs at a multiplicity of infection of 5. Infected cells were harvested 60 h later by centrifugation at 4,000 rpm for 10 min in a Sorvall RC-3B centrifuge. Cell pellets were resuspended in 50 ml of lysis buffer (containing 30 mM HEPES (pH 7.5), 5 mM KCl, and 1 mM MgCl_2_) with 0.1% saturated phenylmethylsulfonyl fluoride and 1 µg/ml each of aprotinin, leupeptin, and pepstatin. Following a 30 minute incubation on ice, the cells were lysed by sonication, the suspension was adjusted to 100 mM KCl, and the lysate was clarified by centrifugation at 20,000 × g for 30 minutes. Forty milliliters of the sample was loaded onto a 5-ml heparin HiTrap column pre-equilibrated with 30 mM HEPES (pH 7.5), 100 mM KCl, 1 mM EDTA, and 10% (v/v) glycerol at a flow rate of 2 ml/min. After washing with the starting buffer, the column was eluted using a 60 ml gradient of KCl (100–450 mM) in 30 mM HEPES-KOH (pH 7.5), 1 mM EDTA, and 10% (v/v) glycerol. hMutLα fractions eluting at approximately 230 mM KCl were diluted to 80 mM KCl using 30 mM HEPES (pH 7.5), 0.1 mM EDTA, and 10% (v/v) glycerol. The diluted sample was then loaded onto a 1-ml Mono Q column pre-equilibrated with buffer A (30 mM HEPES, pH 7.5, 0.1 mM EDTA, and 10% (v/v) glycerol) containing 80 mM KCl at a flow rate of 0.5 ml/min. After washing with starting buffer, the column was eluted with a 20 ml gradient of KCl (80–380 mm) in buffer A. hMutLα fractions eluting at approximately 220 mM KCl were dialyzed against a buffer containing 30 mM HEPES (pH 7.4), 0.1 mM EDTA, 10% (v/v) glycerol, and 150 mM KCl and concentrated with Ultra 0.5 centrifugal filters (MilliporeSigma).

For purification of Flag-tagged MutLα, Sf9 cells expressing Flag-MLH1 and untagged PMS2 were used. Cells were cultured and infected with baculovirus constructs as above. Infected cells were harvested 60 h later by centrifugation at 4,000 rpm for 10 min in a Sorvall RC-3B centrifuge. Frozen cell pellets were resuspended in 30ml lysis buffer with protease inhibitors and lysed initially by sonication. KCl was adjusted to 200 mM using 2M KCl, and further lysed using a 40ml Dounce homogenizer using a tight pestle on ice. The lysate was clarified by centrifugation at 13,500 *x g* for 30 minutes. Binding to M2 Flag-agarose resin (Sigma) was performed at 4°C with gentle rotation. The beads were then washed extensively with lysis buffer containing 200mM KCl, once with TAP wash buffer, and then eluted with TAP buffer containing 0.4 mg/ml Flag peptide (Sigma).

#### *In vitro* pulldown assays

M2 Anti-Flag agarose beads were washed with TAP buffer and 100mM glycine, then blocked with 10% BSA in 1X TAP buffer. To 30 μl of the blocked beads, 2µg of the indicated His-Flag glycosylase was added, along with untagged MutLα (0.3 µg) in a total volume of 100 μl. After rotation at 4°C for one hour, beads were washed five times with TAP wash buffer and once with 1x PBS. The beads were resuspended in 2x Laemmli buffer and incubated at 95°C for five minutes. Samples were analyzed by SDS-PAGE and Coomassie blue staining or western blotting with the indicated antibodies.

#### Immunofluorescence microscopy

All immunofluorescence microscopy was performed as previously described ^72^, with minor modifications. RPE-1 cells were treated with MMS in complete medium at 37 °C for 1h. Next, cells were pulsed with 10µM EdU for 15 min. Cells were pre-extracted with 1× PBS containing 0.2% Triton X-100 and protease inhibitors (ThermoFisher) for 10 min on ice before fixation with 3.5% paraformaldehyde. The cells were then washed extensively with immunofluorescence wash buffer (1× PBS, 0.5% NP-40, and 0.02% NaN_3_), then blocked with immunofluorescence blocking buffer (immunofluorescence wash buffer plus 10% FBS) for at least 30 min. EdU was labeled with AlexaFlour Azide 488 (Thermo Fisher). Primary antibodies were diluted in immunofluorescence blocking buffer for one hour at 4 °C. After staining with secondary antibodies (conjugated with AlexaFluor 488 or 594; Thermo Fisher) and Hoechst 33342 (MilliporeSigma), coverslips were mounted using ProLong Gold antifade reagent (ThermoFisher). Fluorescence microscopy was performed on an Olympus fluorescence microscope (BX-53), using an UPlanS-Apo 100×/1.4 numerical aperture oil immersion lens and cellSens Dimension software. At least 100 cells per sample were analyzed using ImageJ to calculate the total integrated intensity.

For abasic site detection, a doxycycline-inducible HA-APE1 E96Q/D210N plasmid (a gift from Dr. Priyanka Verma, Washington University) was transduced into control RPE-1 cells, or PMS2 or MLH1 knockout cells. In order to test the role of MPG mutants on abasic site production, endogenous MPG was knocked down using lentiviral shRNA, then WT or MIP box mutant MPG constructs (YF-AA and 5D) were expressed via retroviral infection. A negative control sample was included by infecting with an empty virus. APE1 mutant expression was induced with 1ug/ml doxycycline for 6 hours cells were treated with 200-400µM MMS (MilliporeSigma) for one hour. Immunofluorescence was then performed as above.

#### qRT-qPCR

RNA was extracted using the RNeasy Mini Kit (QIAGEN). Reverse transcription was performed on 2μg purified RNA using the High-capacity cDNA reverse transcription kit (Thermo Fisher). PowerUp SYBR Green Master Mix (Thermo Fisher) was used with qPCR using the QuantStudio 6 Flex Real-Time PCR System (Thermo Fisher). Relative quantification was performed using the 2^-ΔΔCt^ method.

#### Colony formation assays

Parental or knockout cells were trypsinized, counted, and plated at low density. After overnight incubation, the cells were treated with the indicated doses of MMS in complete medium. The cells were incubated for 12-14 days, fixed, and stained with crystal violet. The experiment was performed in at least triplicate for each cell line and drug dose. Colonies were counted and relative survival was normalized to untreated controls.

#### *In vitro* glycosylase assays

Glycosylase activity was assessed using a gel-based cleavage assay. Each 10 μL reaction contained reaction buffer (50 mM Tris-HCl pH 7.5, 50 mM KCl, 1 mM DTT, 100 μg/mL BSA, 1 mM MgCl_2_), 80 nM of labeled substrate oligonucleotide, and the indicated amount of glycosylase and MutLα. This substrate was pre-annealed to either a short DNA oligonucleotide to create a gapped structure or to its full complement to form a blunt-ended double-stranded (DS) substrate, as indicated. Reactions were incubated at 37°C for the specified durations and then stopped by adding 2X loading buffer (95% deionized formamide, 0.05 mM EDTA, 1% bromophenol blue). Where indicated, about 0.2 M NaOH was included in the quenching step to induce DNA nicking at the AP sites. The samples were denatured at 90°C for 5 minutes and subsequently resolved on a 12% urea gel at 150 V. The resulting cleavage products were visualized by scanning the gel on a LI-COR Odyssey imaging system. The substrate and product bands were quantified using ImageJ.

#### EMSA

Increased amounts of MPG were incubated with 8 nM of 34 bp HX double-stranded Cy5.5-labeled DNA substrate in reaction buffer (20 mM HEPES, pH 7.5, 100 mM KCl, 1mM MgCl2, 1 mM DTT, 0.25 mg/ml BSA,5% glycerol, and 2 mM ATP) in the presence or absence of MutLα. The mixture was incubated for 20 minutes at room temperature in a final reaction volume of 10 μl. Samples were immediately loaded on 6% polyacrylamide (37.5:1, acrylamide:bis) native gel and run at 100 V for 50 min at 4°C in 0.5X TBE. DNA bands were visualized using LI-COR Odyssey imaging system. The percentage of total DNA bound by each protein was determined by measuring the band intensity present in the bound states and dividing by the total band intensity in the lane. The percentage of DNA bound in each reaction was plotted against the concentration of MPG.

#### Deoxynucleoside quantification by HPLC-MS/MS

RPE-1 cells were treated with the indicated dose of MMS or MNNG for 1 hour, then harvested and processed for DNA purification using the DNeasy Blood and Tissue Kit (Qiagen, 69506). After extraction, 2 μg of genomic DNA was digested with 3 μL of Nucleoside Digestion Mix (NEB, M0649S) in a total volume of 50 μL at 37°C overnight. The digested sample was filtered with a 0.22 μM filter and then processed for nucleoside mass spectrometry analysis ^75,76^.

Deoxyucleoside standards were run in parallel with the samples to verify the chromatographic signals of each deoxynucleoside. Chromatographic separation was performed using an Agilent 1290 Infinity II UHPLC system with a ZORBAX RRHD Eclipse Plus C18 2.1 x 50 mm (1.8 um) column. The mobile phase consisted of water and methanol (with 0.1% formic acid) run at 0.5 ml/min. The run started with a 3 min gradient of 2-8% methanol, followed by a sharp increase to 98% methanol which was maintained for 4 min. Mass spectrometric detection was performed using an Agilent 6470 Triple Quadrupole system operating in positive electrospray ionization mode, monitoring the mass transitions 282/166 (for O6me-dG), 268.1/152.3 (for dG), 266.13/150 (for 1me-dA), 252.11/136 (for dA). The data were visualized with the Agilent Qualitative Navigator analysis software, and the chromatographic peaks of each deoxynucleoside shown with the samples were verified based on the peaks shown with the corresponding standards.

### QUANTIFICATION AND STATISTICAL ANALYSIS

Statistical analyses were performed using Prism 10 (GraphPad Software). Specific tests used for each experiment are described in the figure legends and results section. Data are presented as mean ± SD unless otherwise indicated. Significance was defined as follows: n.s., not significant; \**p* < 0.05; \*\**p* < 0.01; \*\*\**p* < 0.001; \*\*\*\**p* < 0.0001. All experiments were independently repeated at least three times unless stated otherwise. Statistical significance for immunofluorescence data was assessed using the Mann–Whitney test. For in vitro glycosylase assays and EMSA, an unpaired t-test was used. For colony formation assays, two-way ANOVA followed by Bonferroni’s multiple-comparison test was applied.

## Supplemental Figure Legends

**Supplemental Figure S1.**
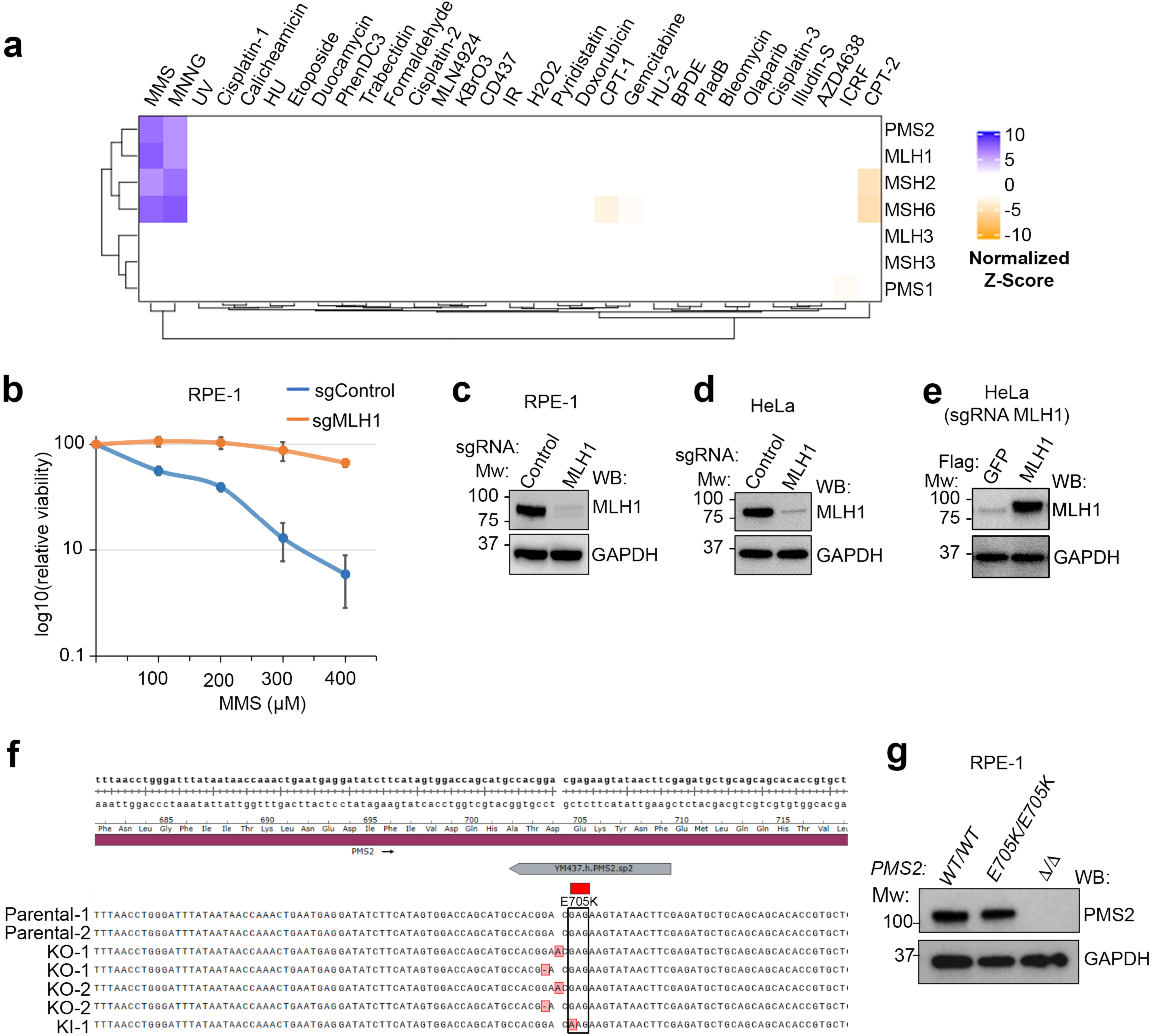
Loss of MMR alters cellular responses to MMS and MNNG. **(a)** Data mining of published CRISPR/Cas9 screening datasets (22) in RPE-1 cells treated with a panel of DNA-damaging agents. Screenshot obtained from https://durocher.shinyapps.io/GenotoxicScreens/ illustrates that knockout of mismatch repair (MMR) genes selectively confers resistance to MMS and MNNG in cells. **(b)** Colony formation assays comparing wild-type (WT) and MLH1 knockout RPE-1 cells following MMS treatment, demonstrating that loss of MLH1 confers increased resistance to MMS. Data represent the mean of three biological replicates ± SD (*p* < 0.0001 as assessed by two-way ANOVA followed by Bonferroni’s multiple-comparison test). **(c)** Western blot analysis confirming MLH1 knockout in RPE-1 cells. **(d)** Western blot analysis confirming MLH1 knockout in Hela cells. **(e)** Western blot analysis confirming Flag-MLH1 expression in MLH1 knockout Hela cells. **(f)** Next-generation sequencing (NGS) validation of PMS2 knockout and homozygous PMS2*^E705K/E705K^* knock-in (KI) mutation. **(g)** Western blot analysis confirmed PMS2 knockout in RPE-1 cells and verified the normal expression of the PMS2 knock-in mutation.

**Supplemental Figure S2.**
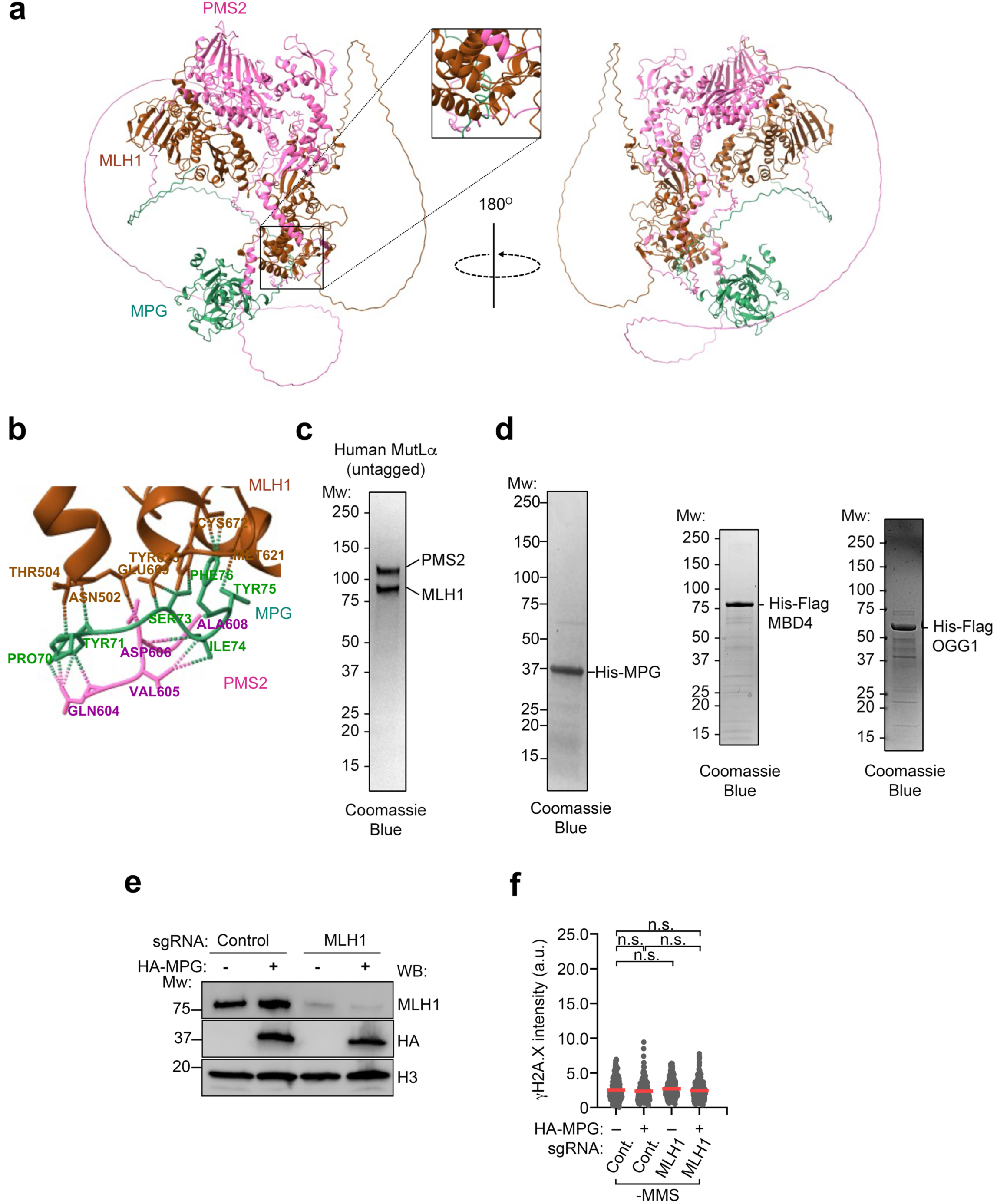
Structural modeling and protein purification of MPG and MutLα. **(a)** Structural modeling performed using AlphaFold3 with MPG (in green) and the two subunits of MutLα (MLH1 and PMS2; in brown and magenta, respectively). **(b)** AlphaFold3 structural analysis of the MPG-MLH1-PMS2 complex highlighting the predicted interacting amino acid residues. **(c-d)** Coomassie Brilliant Blue–stained SDS-PAGE gel purification of recombinant MutLα **(c)**, as well as MPG, MBD4, and OGG1 **(d)**. **(e)** Western blot analysis confirming MLH1 KO and HA-MPG overexpression in RPE-1 cells. **(f)** Immunofluorescence (IF) analysis of γH2AX foci intensity in untreated RPE-1 cells. Statistical significance was determined using the Mann–Whitney test. Data represent the mean of three biological replicates.

**Supplemental Figure S3.**
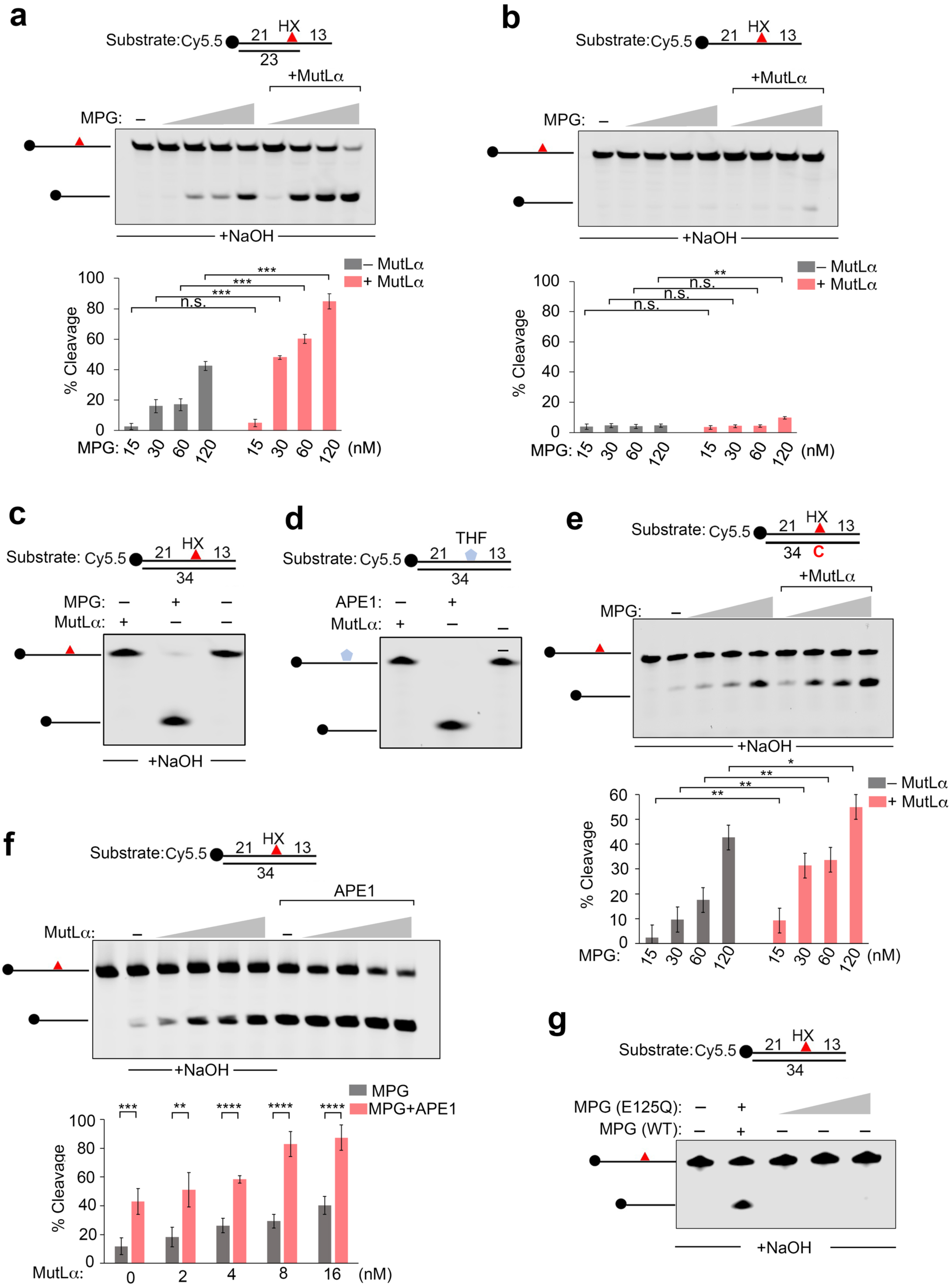
*In vitro* characterization of MPG glycosylase activity and its regulation by MutLα and APE1. **(a-b)** *In vitro* glycosylase assays using hypoxanthine (HX)-containing DNA substrate (80 nM) to evaluate the effect of MutLα (30 nM) on MPG (15-120nM). Assays were performed using either HX substrates annealed with a short complementary oligonucleotide to generate a ssDNA–dsDNA junction **(a)** or ssDNA substrate alone **(b)**. **(c)** *In vitro* glycosylase assays assessing the activity of MutLα (100 nM) alone on duplex HX-containing substrate, with MPG (120 nM) used as a positive control. **(d)** *In vitro* cleavage assays using tetrahydrofuran (THF)-containing DNA substrates to test whether MutLα (100 nM) alone exhibits activity toward abasic-site mimics, with APE1 used as a positive control. **(e)** *In vitro* MPG glycosylase assays using conditions as in Figure S3a-b but using an HX-containing DNA substrate (80 nM) annealed to complementary strand containing “C” opposing the HX. **(f)** *In vitro* MPG glycosylase assays using HX-containing substrate (80 nM) to examine the combined effects of MutLα at the indicated concentration with or without APE1. **(g)** *In vitro* MPG glycosylase assays using HX-containing DNA substrate with wildtype MPG (30 nM) or the catalytically inactive MPG mutant (30, 60, or 120 nM). Data represent the mean of at least three independent replicates ± SD of the mean. Statistical significance was determined using student t-test. Statistical significance was defined as \**p* < 0.05, \*\**p* < 0.01, *** *p* < 0.001, and \*\*\*\**p* < 0.0001.

**Supplemental Figure S4.**
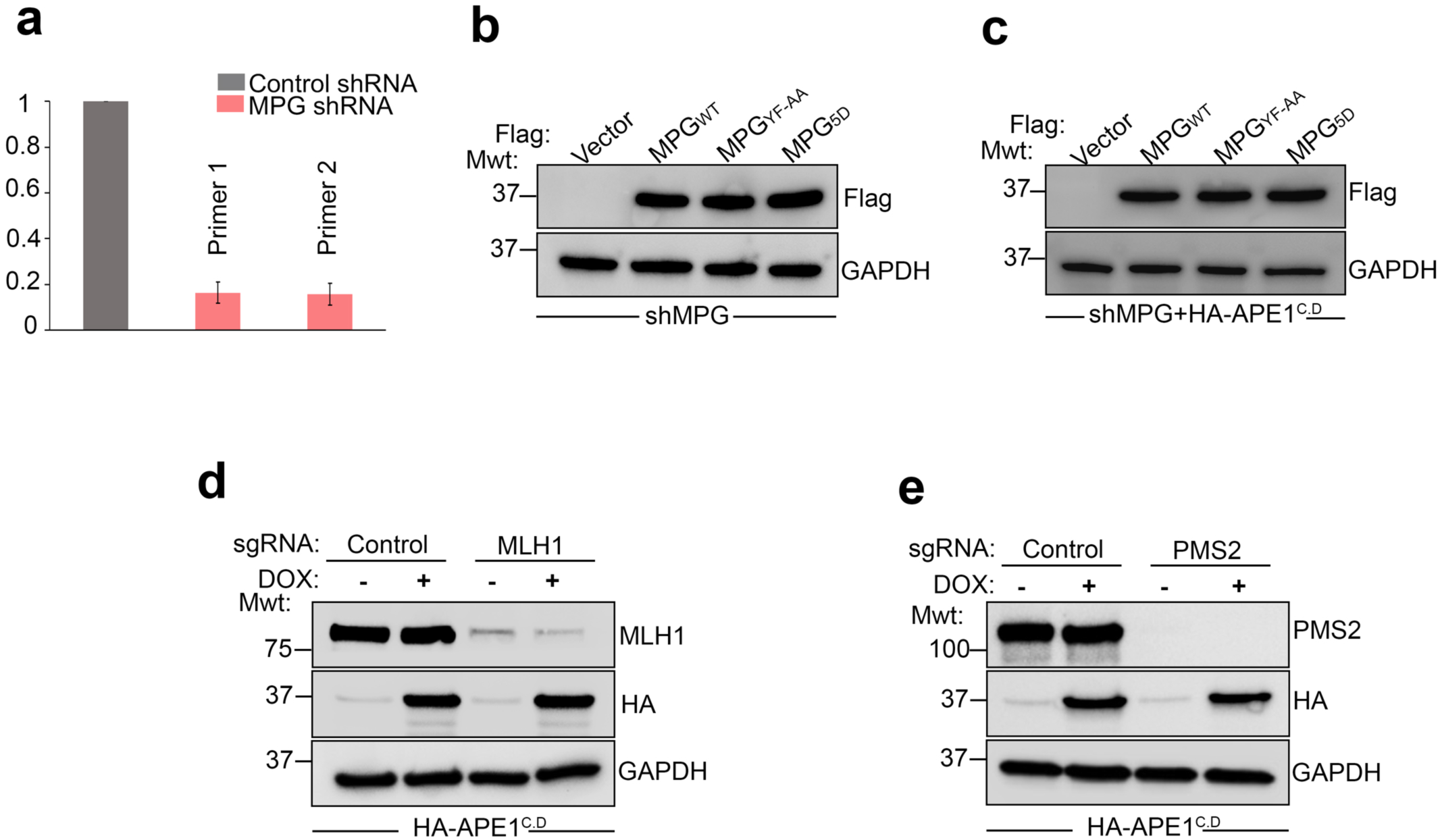
Validation of MPG knockdown, overexpression, and MLH1 genetic manipulations. **(a)** Quantitative RT-PCR (qRT-PCR) analysis confirming knockdown of endogenous MPG expression, with normalization against β-actin. **(b-c)** Western blot analysis of Flag-MPG and MPG mutants as indicated in RPE-1 cells. **(d)** Western blot analysis of RPE-1 cells expressing control or MLH1-targeting sgRNA, with or without doxycycline-induced expression of HA-tagged catalytically inactive APE1. **(e)** Western blot analysis of control or PMS2-KO RPE-1 cells with or without doxycycline-induced expression of HA-tagged catalytically inactive APE1.

